# SpliceViNCI: Visualizing the splicing of non-canonical introns through recurrent neural networks

**DOI:** 10.1101/2020.02.09.940551

**Authors:** Aparajita Dutta, Kusum Kumari Singh, Ashish Anand

## Abstract

Most of the current computational models for splice junction prediction are based on the identification of canonical splice junctions. However, it is observed that the junctions lacking the consensus dimers *GT* and *AG* also undergo splicing. Identification of such splice junctions, called the non-canonical splice junctions, is also essentially important for a comprehensive understanding of the splicing phenomenon. This work focuses on the identification of non-canonical splice junctions through the application of a bidirectional long short-term memory (BLSTM) network. Furthermore, we apply a back-propagation based (integrated gradient) and a perturbation based (occlusion) visualization techniques to extract the non-canonical splicing features learned by the model. The features obtained are validated with the existing knowledge from the literature. Integrated gradient extracts features that comprise contiguous nucleotides, whereas occlusion extracts features that are individual nucleotides distributed across the sequence.

## 1. Introduction

Alternative splicing (*AS*), a splicing mechanism, allows a pre-mRNA to produce distinct mature mRNA. *AS* does so by differentially joining or skipping exons and introns of a gene.[1] *AS* contributes to proteome diversity. It is estimated that about 92-94% of multi-exonic human genes undergo *AS*.[2] Furthermore, exonic mutations perturb *AS* [3] and about 1/3 of disease-associated alleles alter splicing.[4] Thus, comprehensive understanding of *AS* is important in our endeavour of understanding complex transcription mechanism.

Majority of the studies, especially computational studies, focus on studying splicing events associated with highly conserved consensus dimers *GT* and *AG* at the exon-intron (*donor site*) and intron-exon (*acceptor site*) boundaries.[5] Donor and acceptor sites are commonly referred as splice junctions or sites. Splice sites with the conserved dimers are referred as *canonical splice sites*. However, splice sites lacking the consensus dimers *GT* and *AG* (*non-canonical splice sites*) at the junctions have been observed. Such observations suggest distinct regulation pathways in alternative splicing.

The importance of unraveling the hidden biological features of non-canonical splicing is manifold. Understanding non-canonical splicing events can help explain the unconventional regulation pathways for which a function has not yet been identified.[6] Non-canonical splicing most often has a role in regulating gene expression.[7] Mutations in the genomic regions far away from canonical splice sites can cause disease by disrupting the splicing mechanism and activating non-canonical splice sites via non-canonical splicing. Finally, non-canonical splicing mechanisms can be targeted or exploited for new therapeutic strategies. Such strategies have already been applied to diseases like cancer, muscular dystrophies, and haemophilia.[8, 9]

Our knowledge of non-canonical splice sites and regulatory signaling mechanisms is limited. The limited knowledge hampers the use of computational learning models relying on hand-engineered features. However, the availability of abundant sequence data and the recent advancements in representation learning capability of deep learning models have produced promising results for several genome sequence-based tasks.[10, 11]

Several deep learning models have been applied for the identification of splice junctions as well. However, similar to experimental studies, the majority of these models focus on the identification of canonical splice junctions [12, 13, 14, 15]. The models which include non-canonical splice junctions [16, 17, 18, 19] consider only a limited splicing context at the junctions. The primary objectives of this paper are to analyze the model in the context of extended flanking regions for non-canonical splice junctions and extract the biologically relevant sequence features. Towards these objectives, two ways to generate negative data, and their influence on the model’s performance were also analyzed. In particular, we focus on the following research questions:

- How well can a representation learning model encode non-canonical splicing context?
- What meaningful and biologically relevant features can be extracted from such models?
- What can be said about the extracted features in contrast to the known knowledge about the non-canonical sites?

Towards answering the research questions mentioned above, this paper introduces SpliceViNCI, a model that identifies splice junctions by applying bidirectional long short-term memory (BLSTM) networks. Similar models have already been applied by some of the previous research works to identify splice junctions. [14, 19]. Our research endeavor differs from the previous applications on the following factors.

The first factor is the extraction of biologically relevant features learned by the model for both canonical and non-canonical splice junctions. Most of the previous studies focused only on the canonical splice junctions. Byunghan et al.[14] did consider the prediction of non-canonical splice junctions as well but did not extract any biologically relevant features. However, our study extensively and systematically analyzes the non-canonical splicing phenomenon through visualization of the features learned by the model. Our previous work, namely SpliceVisuL[19], did focus on identification and feature extraction of non-canonical splice junctions as well but considered a limited flanking region of 40 nt.

Furthermore, Dutta et al.[19] inferred the presence of subtle non-canonical splicing features beyond the 40 nt context. The inference was based on the combination of the results from SpliceVisuL analysis and the existing knowledge. SpliceVisuL analysis indicated that non-canonical splicing signals are in the form of isolated nucleotides rather than consecutive motifs. This was further supported by the existing knowledge that non-canonical splicing signals partially lack the known consensus, and often the signals are scattered far from the splice junctions.[20] SpliceViNCI explores the hypothesis of the presence of signals in an extended region by considering a larger context in the vicinity of splice junctions.

SpliceViNCI targets to attain optimal performance for the identification of canonical as well as non-canonical splice junctions. We extract the non-canonical splicing features learned by the model and validate them with the existing knowledge. Furthermore, the attention layer present in SpliceVisuL is removed from SpliceViNCI since the attention layer failed to capture all the relevant features in SpliceVisuL.[19] It is also worth noting that, unlike spliceVisuL, SpliceViNCI is trained with the negative dataset that comprises both canonical and non-canonical splice junctions. This is done so that the model can recognize the more subtle non-canonical splicing signals apart from the signals at the junctions.

The contributions of this work can be summarized as follows:

- We present a BLSTM model named SpliceViNCI, which attains state-of-the-art performance for the identification of both canonical and non-canonical splice junctions.
- We design two datasets, Type-1 and Type-2, based on two different sampling strategies to generate negative data.
- We analyze the performance of various state-of-the-art models as well as the proposed model with both the datasets.
- We analyze the length of the flanking region required for obtaining the optimal performance in the identification of non-canonical splice junctions.
- We apply two effective visualization techniques to discern the non-canonical splicing features learned by the model. The findings thereof are validated with the existing knowledge from the literature.

## 2. Background

The advent of advanced sequencing technologies like RNA-seq has produced a plethora of sequenced data, which boosted both alignment-based and machine learning-based methodologies for identifying splice junctions. However, the existing alignment-based methods [21, 22] mostly identify only the canonical splice junctions.[16] The traditional machine learning-based methods rely on hand-engineered features extracted from the sequence neighboring the splice junctions for splice junction identification.[23, 24, 25, 26, 27] The limited knowledge of the splicing features may lead to the inclusion of irrelevant features that can adversely affect the model’s performance. Several feature selection methods have been applied to obtain the optimal feature set.[28, 29] However, the potential of these methods is still limited by the existing knowledge bias.

The emergence of deep learning era led to the application of models that learn the complex splicing signals from the genome sequence *de-novo*. There are implementations of deep Boltzmann machine [16], deep convolutional neural networks (CNN) [17, 12, 13], distributed representation learning [18], recurrent neural networks (RNN) [14, 19], and deep residual neural network [15] for the prediction of splice junctions. Some of these models are based on the identification of only canonical splice junctions. The models which include non-canonical splice junctions [16, 17, 18, 19] consider limited splicing context at the junctions. These methods extract upstream and downstream sequences, at both donor and acceptor junctions, of a length ranging from 30 to 40 nucleotides (nt) since this length is considered as optimal for splicing signals in various studies.[23, 30] Also, these models target on achieving the overall optimal performance considering both canonical and non-canonical splice junctions together.

A major limitation of the deep learning models is their inability to explain the logistics behind their predictive performance due to their ‘black box’ nature. To alleviate this shortcoming, several visualization techniques are applied to extract the features learned by the model. Most of the widely used visualization techniques can be categorized as perturbation based and back-propagation based techniques.[31] Back-propagation based visualization technique called *DeepLIFT* [32] has been applied in the extraction of canonical features learned by a CNN.[13] Both perturbation based and back-propagation based techniques have been applied in the extraction of canonical and non-canonical features learned by an RNN.[19] However, the research effort by Dutta et al. [19] revealed the existence of non-canonical splicing features far beyond the genomic context considered in the work.

## 3. Methods

This section elaborates on the neural network architecture employed for the classification of true and decoy splice junctions. We subsequently describe the visualization techniques applied to extract the relevant features learned by the model.

### 3.1. Neural architecture

The overview of the network architecture is shown in Figure 1.

**Figure 1:**
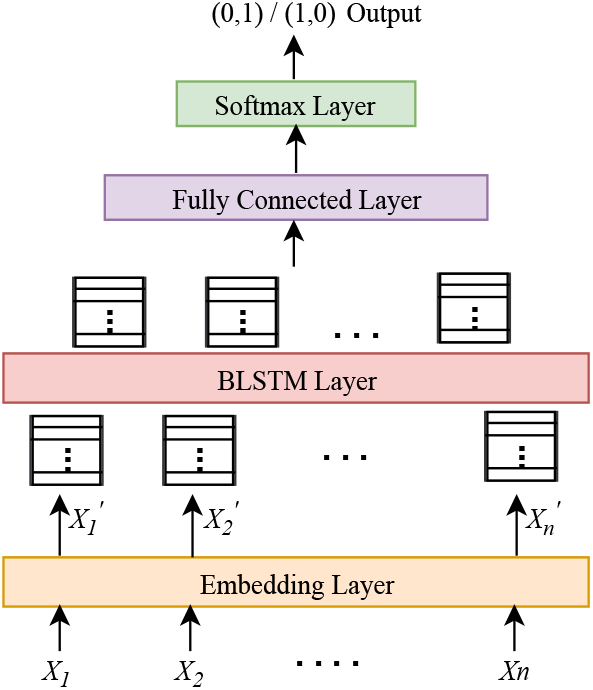
A schematic of the network architecture.

#### 3.1.1. Input representation

The input to the learning model is the genome sequences extracted from the vicinity of splice junctions. These sequences comprise the four nucleotides: *A* (*Adenine*), *C* (*Cytosine*), *G* (*Guanine*), *T* (*Thymine*) and *N* (*denoting any one of the four nucleotides*). The input sequences are passed through an embedding layer to generate a *k*-dimensional dense representation for each of the five nucleotide codes. Therefore, an input sequence of length *n* will be transformed into an *n × k* dimensional dense vector that gets updated while training the network. The dense vector is observed to perform better than the traditional one-hot encoded vector due to the learning of meaningful representation through training.[14]

#### 3.1.2. Splice junction representation using BLSTM network

The BLSTM layer comprises the heart of SpliceViNCI, which captures relevant features from the input sequences. The BLSTM layer is formed by two identical hidden layers composed of LSTM units. LSTM units are like memory cells capable of learning long-term dependencies.[33, 34] There have been several refinements in the structure of an LSTM unit.[33, 35, 36] We apply the LSTM structure proposed in Gers et al. [34]. This LSTM unit contains input (*i*_*t*_), forget (*f*_*t*_) and output (*o*_*t*_) gates that regulate the information flowing through cell state (*C*_*t*_) at any time step ′*t*′. The information flowing out of the cell at a time step ′*t*′ for input *X*_*t*_ is denoted by the hidden value *h*_*t*_.

The activation vectors *i*_*t*_, *o*_*t*_, *f*_*t*_, and 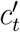, for the input gate, output gate, forget gate and candidate cell state at time step *t*, can be represented by the following equations:

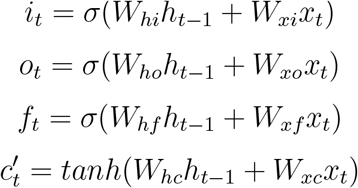

The actual cell state value (*C*_*t*_) at time step *t* is computed by:

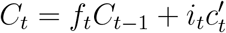

Finally, the output (*h*_*t*_) of the unit at time step *t* is given by:

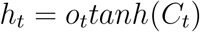

The two LSTM layers process the input sequence in forward and backward directions simultaneously. The output *h*_*t*_ of the BLSTM layer is given by the concatenation of the output generated by the two LSTM layers. This can be written as

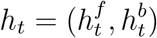

where 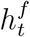 and 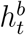 are the output generated in the *t*^*th*^ time step by the forward and backward LSTM layers, respectively. The LSTM units capture dependencies among input subsequences as it processes the input. Hence, 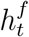can be expected to capture the relationship of an input subsequence with the subsequence to its left. Likewise, 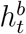 can capture the relationship of an input subsequence with the subsequence to its right. Thus, the concatenated output *h*_*t*_ is a more comprehensive feature representation of an input splice junction.

The *n* × *k* dimensional dense vectors generated from the embedding layer are fed into the BLSTM layer. Each LSTM layer generates an *n* × *n*_*l*_ vector which is concatenated to form an *n* × 2*n*_*l*_ vector as the output from BLSTM layer. The output from the BLSTM layer is fed into a fully connected layer and finally to the output layer, which classifies the input sequence as a true or decoy splice junction. Binary cross-entropy and Adam [37] are the loss function and the optimizer applied.

### 3.2. Visualization techniques

We employ two effective visualization techniques, namely integrated gradients and occlusion, for the extraction of relevant splicing features learned by the model. Both the visualization techniques assign an importance value to each nucleotide position in a genome sequence. We name the importance value as *deviation value*. Higher *deviation value* for a sequence position implies that position as more significant in the identification of the splice junction. The visualization techniques are further explained in the subsequent section.

#### 3.2.1. Integrated gradients

Integrated gradients is a back-propagation based visualization technique proposed by Sundararajan et al.[38]. This technique involves computing the gradient of the output stepwise along the linear path from a chosen baseline to the given input. For sequence-based models, the baseline is usually the zero embedding vector. The number of steps is a hyperparameter, which is chosen as 50 for our experiments. The integrated gradient is computed by averaging the gradients obtained from each step along the linear path. This average gradient at each sequence position is the *deviation value* in this case.

The reason for the selection of integrated gradients over other back-propagation based visualization techniques like basic gradients, DeepLIFT [32], and Layer-wise relevance propagation (LRP) [39] can be explained by the following factors. The integrated gradient satisfies two desirable properties of attribution methods: sensitivity and implementation invariance, either of which is not satisfied by basic gradients, DeepLIFT and LRP.[38]

An attribution method is said to satisfy the sensitivity property when for any input-baseline pair, which differs in one feature and has different predictions, the differing feature is assigned a non-zero attribution. This property is desired in any attribution method because the lack of this property intuitively means that the attribution method may focus on irrelevant features. Gradient based methods lack this property, whereas methods like DeepLIFT and LRP satisfy it.[38]

However, DeepLIFT and LRP lack the property of implementation invariance. An attribution method is considered invariant to the implementation model when it assigns identical feature attributions for functionally equivalent models. Lack of this property may lead to the attribution methods being sensitive to the unimportant biases of the implementation model.[38] Also, in models like RNN with multiplicative interactions (like LSTM and BLSTM), DeepLIFT fails to produce meaningful results.[40] Although there are researches inferring that a more principled backpropagation rule can produce improved results for DeepLIFT in the case of multiplicative units.[41, 42]

#### 3.2.2. Occlusion

Occlusion is a perturbation based visualization technique proposed by Zeiler et al..[43] As the name suggests, this method obstructs or occludes a portion of the input by masking it. We occlude an input sequence by moving a sliding window of length one, along the sequence, replacing one nucleotide each time with *N*. The modified sequence is passed through the trained model to obtain the difference between the output with and without occlusion. The difference between the two output values with and without occlusion of *n*^*th*^ sequence position gives the *deviation value* of that position in the sequence.

For one input sequence of length *L*, we obtain *L* masked inputs, each one with a different masked position. Hence, for a batch of *B* sequences, we have a *B* × *L* input matrix. For a computationally efficient implementation of this technique, we prepare all the *L × L* masked sequences for a given input sequence beforehand. Eventually, for a batch of size *B*, we obtain (*B* * *L*) × *L* input matrix, which is fed into the model in a single batch. We obtain (*B* * *L*) ×1 output matrix, which is reshaped to obtain the required *B* × *L* matrix of deviation values. We propose two variations of occlusion, as discussed next.

##### Fixed length occlusion

In this type of occlusion, we consider a fixed window length of *k* nucleotides for masking the input. This is denoted by *occlusion-k*. The *deviation value* of the window is obtained by summing up the *deviation values* of each nucleotide position within the window. This summed up *deviation value* is added to the position (*k* + 1)*/*2 lying in the center of the window.

##### Variable length occlusion

In fixed length occlusion, we consider only a fixed length genomic region for detecting the splicing features, whereas, in a real scenario, the relevant features may be of variable length. Hence, in this variation of occlusion, we consider different occlusion windows *w*_*l*_ of length *l* ∈{ 1, 3, 5, 7, 9, 11 } at each sequence position. The *deviation value* for each window length is computed, and the highest deviation value is chosen for that sequence position. The corresponding window length is saved in another matrix called the *window matrix*. Therefore, the value in *j*^*th*^ column of *i*^*th*^ row of the *window matrix* denotes the length of the pattern, centered at position *j* of the *i*^*th*^ sequence, that contributes maximum to the prediction of the model.

## 4. Experimental Setup

### 4.1. Dataset

The procedure of generating the positive and negative data is described in the following subsection.

### 4.1.1. Positive data

We assess the performance of SpliceViNCI on the identification of novel splice junctions. We, therefore, generate the training and test dataset from two different versions of GENCODE [44] annotations based on human genome version *GRCh38*. Each sample in the data is an intron that comprises donor-acceptor junction pair. The introns are extracted from protein-coding genes only.

290,502 junction pairs are extracted from version 20 as the training data, whereas 293,889 splice junctions are extracted from version 26. The test data is composed of only those introns which were not annotated in version 20. This yields 5,612 novel junction pairs in the test data. We consider introns of length greater than 30 nt only since an existing study [45] suggests that introns of length less than 30 nt can be attributed to sequencing errors. Further, each intronic sequence is truncated to a fixed length by chipping and concatenating the donor and acceptor junctions with a certain length of flanking region (see Section 5.2).

#### 4.1.2. Negative data

Based on the type of features captured in the data, two variants of negative data is generated. Both randomness-based and consensus-based negative data are described in the following section.

##### Randomness-based negative data

We extract a subsequence from the center of an intron with the safe assumption that no splice junction will be present between a pair of donor-acceptor junctions. This procedure of negative data generation is proposed by Noordewier et al..[46] The lengths of positive data and the extracted subsequence are taken equal. The non-randomness in genome sequences is captured in this type of negative data. We obtain 290, 502 false samples for training data and 5, 612 false samples for test data using this procedure.

##### Consensus-based negative data

Since we are seeking a deeper insight into non-canonical splicing, therefore, it is necessary to mimic similar sequences in the positive and negative data, so the model learns to identify the signals regulating non-canonical splicing in particular. Existing study says that more than 98% of splice junctions are canonical, comprising the *GT* and *AG* consensus dimers.[47] The genome sequences have frequent occurrences of the consensus dimers, but not all of those are identified as splice junctions by the splicing machinery. This indicates the presence of other splicing signals that govern the selection of splice sites. Apart from the canonical consensus dimers, there are two other commonly known classes of non-canonical splice junctions, namely the *GC*-*AG* and *AT*-*AC* junction pairs.[48]

Hence, we compose this type of negative data considering randomly selected *GT*-*AG, GC*-*AG*, and *AT*-*AC* dimer pairs in the genome sequence such that the dimers are not actual splice junctions. The idea of training the splice site prediction models with a consensus-based negative dataset has been applied in previous works. [49, 16, 13] These works have used datasets like *NN*269 [49] and *GWH* [27, 16, 13] where the sequences in the negative data comprise only *GT* and *AG* dimers at the donor and acceptor splice junctions, respectively. We added two commonly known non-canonical consensus pairs *GC*-*AG* and *AT*-*AC* to let the model learn features governing non-canonical splicing as well.

We randomly search for the donor site consensus in the genome sequence, followed by the corresponding acceptor site consensus. We name such junctions as the negative splice junctions. The randomly sampled dimers should both lie in the same chromosome. The length of the flanking region is considered equal to that in the positive data. We randomly sample 290, 502 training and 5, 612 test data from the human genome assembly version *GRCh38* using this method. The frequencies of both the classes of non-canonical consensus are considered as 0.5% based on their frequencies reported by Stephen M. Mount [48]. The remaining negative data comprises the canonical *GT* -*AG* dimer pair.

The distribution of canonical and non-canonical splice junctions in the positive and negative data of both training and test dataset is shown in Table 1. We form two types of dataset: Type-1 and Type-2. The Type-1 dataset comprises the positive data and randomness-based negative data, whereas the Type-2 dataset comprises the positive data and consensus-based negative data.

**Table 1:** Distribution of canonical and non-canonical splice junctions in the positive and negative data.

| Dataset | # of training samples |  | # of test samples |  |
| --- | --- | --- | --- | --- |
|  | canonical | Non-canonical | canonical | Non-canonical |
| Positive | 255674 | 5777 | 5241 | 371 |
| Randomness-based negative | 1096 | 260355 | 14 | 5598 |
| Consensus-based negative | 258939 | 2512 | 5557 | 55 |

The distribution of the two most frequent non-canonical dimer pairs in the training and test dataset is shown in Table 2. In the positive training data, we see that the two most frequent non-canonical dimer pairs conform to the non-canonical dimers reported by Stephen M. Mount [48]. The frequency depicts the count and percentage of non-canonical sequences comprising a particular dimer-pair. As expected, the randomness-based negative data has a smoother frequency distribution of dimer-pairs since it is not governed by any biases and, therefore, only represents the randomness of a genome sequence.

**Table 2:** Distribution of the top 2 most frequent non-canonical dimer pairs in the positive and negative data.

| Dataset | Training data |  | Test data |  |
| --- | --- | --- | --- | --- |
|  | Top 2 | Frequency | Top 2 | Frequency |
| Positive | GC-AG | 3848 (66.61%) | GC-AG | 165 (44.47%) |
|  | AT-AC | 232 (4.02%) | GA-AG | 16 (4.31%) |
| Randomness-based negative | TT-TT | 2461 (0.94%) | TT-TT | 54 (0.96%) |
|  | AA-TT | 2013 (0.77%) | TG-TT | 47 (0.84%) |
| Consensus-based negative | GC-AG | 1259 (50.12%) | GC-AG | 28 (50.90%) |
|  | AT-AC | 1253 (49.88%) | AT-AC | 27 (49.10%) |

### 4.2. Training and hyperparameter tuning

The training data is partitioned into 90% train and 10% validation data for tuning the hyperparameters of SpliceViNCI. Each nucleotide in the input sequence of length *N* is converted to a 100-dimensional vector by the embedding layer to form an *N* × 100 dense vector. This dense vector is passed through the BLSTM, fully connected, and softmax output layer with 100, 1024, and 2 units, respectively. The values for dropout and recurrent dropout are tuned to 0.5 and 0.2, respectively. We train the model for 10 epochs with a batch size of 128.

### 4.3. Baselines

Our choice of baselines is based on the existing state-of-the-art models in the task of splice site prediction. SpliceRover[13] is a CNN based state-of-the-art model that predicts splice sites and identifies the relevant biological features. The objectives of SpliceRover aligns with the objectives of SpliceViNCI. We have also chosen another CNN based model, namely DeepSplice[17], as a baseline since both DeepSplice and SpliceViNCI formulate the input data in the form of donor-acceptor junction pair. Therefore, it is interesting to compare the performance of both these models for the same task.

Apart from these, we replaced the BLSTM units of SpliceViNCI with LSTM units and considered the LSTM-base model as a baseline. This performance comparison can justify the choice of BLSTM units as the hidden layer of the model. Finally, we choose SpliceVec-MLP[18] as a baseline because of its promising performance over CNN based model. However, SpliceVec-MLP has a limitation of not being able to extract the biological features governing the model performance. We have not considered SpliceVisuL[19] as a baseline since the model in SpliceVisuL is similar to that of SpliceViNCI, except that the attention layer is removed in SpliceViNCI. The attention layer is removed because the layer failed to recognize meaningful features in SpliceVisuL.

The following models are implemented as baselines and the hyperparameters are tuned using the procedure mentioned in Section 4.2. The baselines are trained and tested on the dataset explained in Section 4.1. The tuned hyperparameters and the baseline architectures are as follows:

1. LSTM-base: We replace the BLSTM units in SpliceViNCI with LSTM units. All hyperparameters are the same as that of SpliceViNCI.
2. SpliceRover: This model is a deep CNN proposed by Zuallaert et al.[13] The model identifies acceptor (donor) splice junctions in an acceptor (donor) classification model. The authors propose the use of a different number of convolutional layers for different sequence lengths. We consider two convolutional layers, followed by a max pooling layer based on the optimal performance obtained on our dataset. Tuned values of batch size, learning rate, decay rate, number of steps, and Nesterov momentum are 64, 0.05, 0.5, 5, and 0.9, respectively.
3. SpliceVec-MLP: This model is proposed by Dutta et al.[18] The model generates distributed representations of true and decoy splice junctions using a shallow neural network, which is then classified by a multilayer perceptron.[18] The batch size and learning rate of Adam optimizer are considered as 128 and 0.001.
4. DeepSplice: Zhang et al. proposed this model.[17] This is a deep CNN model that identifies a true donor-acceptor junction pair from a decoy junction pair sequence. The values for batch size, epochs, and Adam optimizer learning rate are tuned to 160, 30, and 0.001, respectively.

## 5. Results

### 5.1. SpliceViNCI learns better representations of non-canonical splice junctions

The quality of the embeddings obtained from the fully connected layer of SpliceViNCI is evaluated by projecting the 1024 dimensional dense vectors into 2 dimensional space using t-SNE.[50] We plot the positive and negative non-canonical test sequences from Type-1 and Type-2 dataset in Figures 2(a) and 2(b), respectively. The points in blue are positive non-canonical sequences, whereas the points in red are negative non-canonical sequences. Similarly, we also plot the embeddings obtained from fully connected layer of SpliceRover for both Type-1 and Type-2 dataset in Figures 2(c) and 2(d).

**Figure 2:**
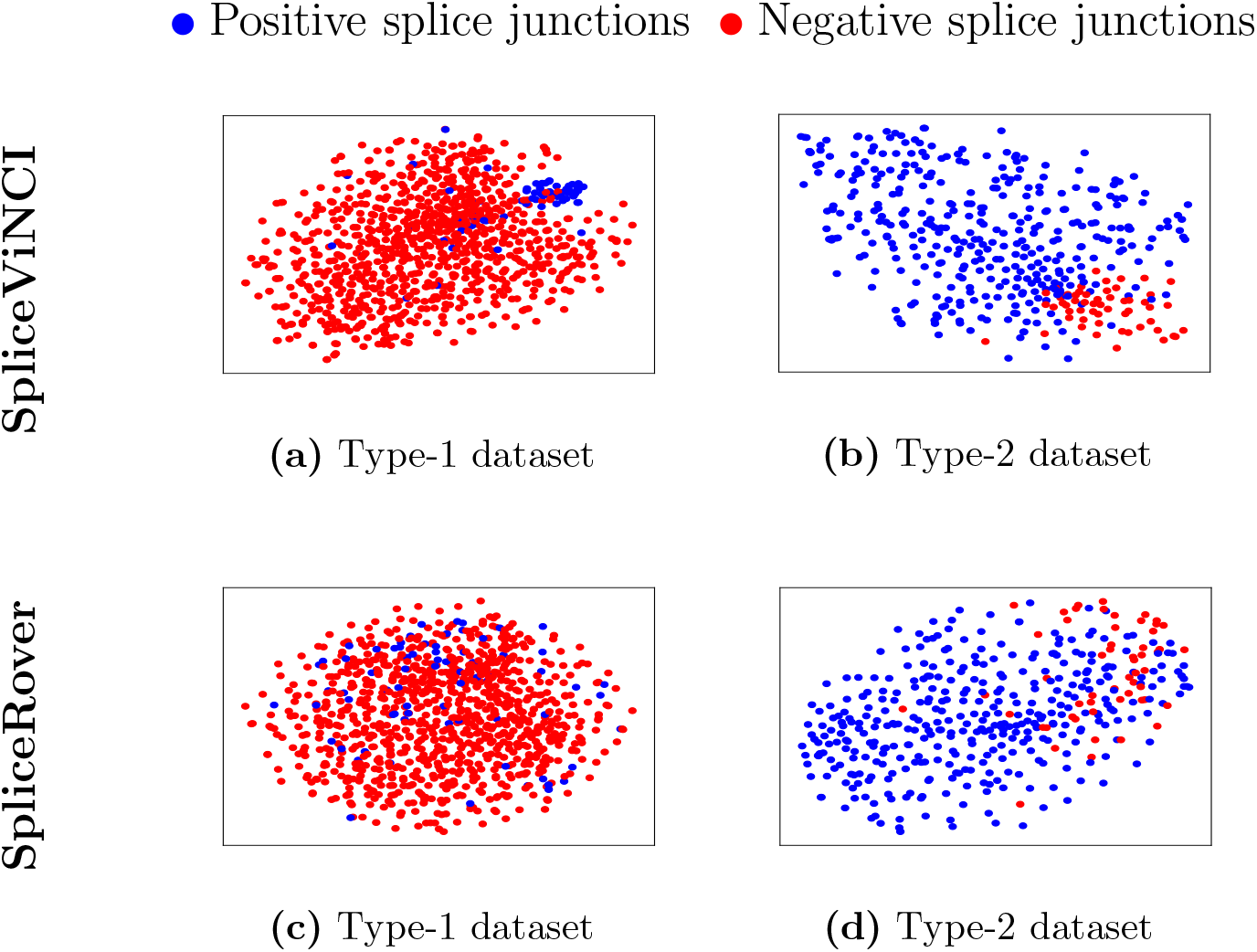
t-SNE plots of non-canonical splice junctions obtained from SpliceViNCI for (a) Type-1 dataset, (b) Type-2 dataset and from SpliceRover for (c) Type-1 dataset, (d) Type-2 dataset. The points in blue are positive splice junctions whereas points in red are negative splice junctions.

Relatively more distinct clusters are observed for both the positive and negative datasets in the case of SpliceViNCI compared to SpliceRover. Since no user-defined features are fed into the model and both the models learn the relevant features *de-novo* from the genome sequences, we can infer that the proposed architecture extracts the splicing features better than SpliceRover in the case of both positive and negative non-canonical splice junctions.

### 5.2. Non-canonical splicing features are relatively further from the splice junctions

Since the length of the flanking region containing important splicing signals is not known, we vary this length in the input sequences to find the optimal flanking region that produces the best performance in splice junction prediction. We vary the flanking region from 20 to 180 nt with a step size of 20 nt. An input sequence comprises upstream and downstream regions of donor and acceptor junctions concatenated in order. Each junction comprises a canonical or non-canonical dimer. Therefore a flanking region of length *N* results in an input sequence of length 4 × *N* + 4.

Table 3 shows the performance of SpliceViNCI in terms of F1-score with a varying flanking region on Type-1 and Type-2 dataset. We obtain the performance of SpliceViNCI on canonical (*can*) and non-canonical (*non-can*) splice junctions separately. This is because the splicing signals may be differently distributed for canonical and non-canonical splicing resulting in different optimal flanking regions in both the cases.

**Table 3:** F1-score (in percentage) obtained by SpliceViNCI in identification of canonical (*can*) and non-canonical (*non-can*) splice junctions with varying flanking region on Type-1 and Type-2 dataset.

| Flanking region | Type-1 dataset |  | Type-2 dataset |  |
| --- | --- | --- | --- | --- |
|  | can | non-can | can | non-can |
| 180 | 99.50 | 69.71 | 99.13 | 97.38 |
| 160 | 99.39 | 65.47 | 99.06 | 97.24 |
| 140 | 99.65 | 72.09 | 99.05 | 97.66 |
| 120 | 99.65 | **74.04** | 99.05 | **97.67** |
| 100 | 99.67 | 73.56 | 99.04 | 96.82 |
| 80 | **99.70** | 70.31 | 99.02 | 96.81 |
| 60 | 99.65 | 71.06 | **99.07** | 96.40 |
| 40 | 99.60 | 69.32 | 98.30 | 95.65 |
| 20 | 98.60 | 60.01 | 95.82 | 93.65 |

We see that the performance of SpliceViNCI in the prediction of non-canonical splice junctions improves by about 14% for Type-1 dataset and 4% for Type-2 dataset with an increase in the context. We also observe that SpliceViNCI obtains comparable performance for both the dataset in case of canonical splice junctions. However, in the case of non-canonical splice junctions, the performance improves significantly in the Type-2 dataset.

SpliceViNCI obtains the maximum F1-score of 97.67% (74.04%) in the Type-2 (Type-1) dataset for the prediction of non-canonical splice junctions. The improvement in performance for the Type-2 dataset can be attributed to the consensus-based negative data in the Type-2 dataset. Since the consensus-based negative data is composed of the splice junction consensuses GT-AG, GC-AG, and AT-AC, the positive and negative data in the Type-2 dataset look very similar at the splice junction. This enables the model to recognize other subtle features in the flanking region apart from the dimers at the splice junctions, which differentiate the true and decoy splice sites. This hypothesis corroborates the rationale behind the similar formulation of negative datasets by Bretschneider et al.[51]

On the contrary, the negative data in the Type-1 dataset comprises sequences from the center of each intron. This type of negative data lacks the consensus dimers and only captures the non-randomness of genome sequences. Therefore, the model recognizes the consensus dimers as the primary features in this type of dataset and may miss out on the other subtle splicing signals that govern non-canonical splicing.

We obtain the optimal performance for the prediction of canonical splice junctions at 60 to 80 nt context. An optimal length of 30 to 40 nt is suggested in the literature [23, 30] and validated in various studies [16, 17, 12, 18, 19] for canonical splice junction prediction. These studies obtained negligible improvement in the model’s performance on further increase of the flanking context.

We obtain the optimal performance for the prediction of non-canonical splice junctions at a flanking region of 120 nt. To examine the statistical significance of the context length, we performed the student’s *t*-test. We formed two groups, each comprising F1-scores obtained from five different executions, with a flanking region of 120 nt and 80 nt, respectively. The *P* -values obtained for Type-1 and Type-2 dataset are 0.002 and 0.003, respectively. We consider a *P*-value < 0.05 as statistically significant.

The statistical significance of variation in the flanking region indicates the presence of non-canonical splicing signals further away from the splice junction. This inference can be validated by the study, which suggests that non-canonical splice junctions may lack some known consensus, or the splicing signals may be distally located from the splice junctions.[20]

### 5.3. SpliceViNCI outperforms state-of-the-art splice junction prediction models

We compute the prediction performance of various state-of-the-art models on Type-1 and Type-2 dataset, considering the optimal flanking region of 120 nt obtained in Section 5.2. F1-score is considered as the performance metric. We see that SpliceViNCI outperforms all state-of-the-art models in the prediction of non-canonical splice junctions in both Type-1 and Type-2 dataset (Figure 3).

**Figure 3:**
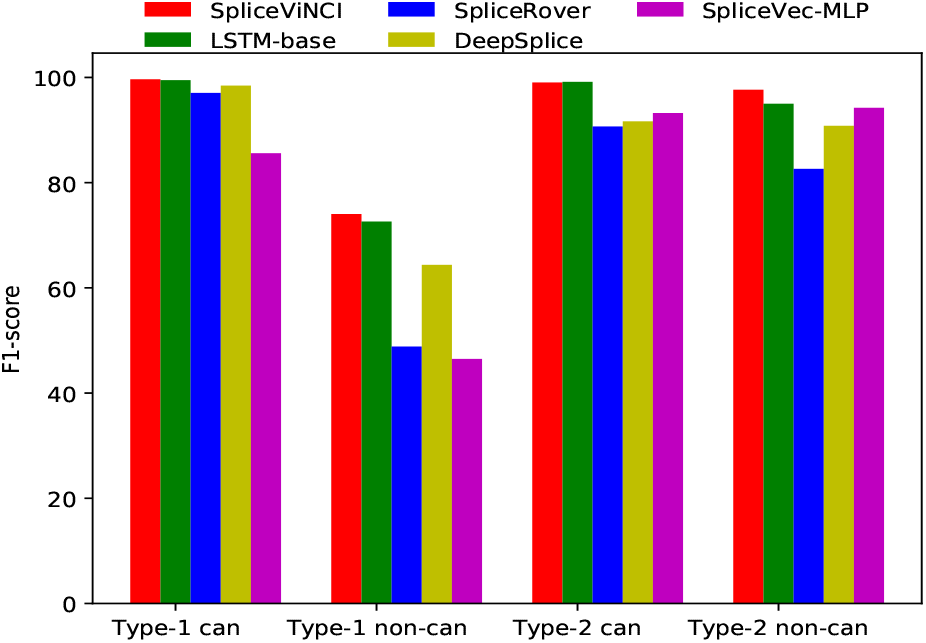
Performance of various state-of-the-art models. The performance is measured in terms of F1-score for canonical (*can*) and non-canonical (*non can*) splice junctions from both Type-1 and Type-2 dataset. F1-score is computed in percentage.

In the prediction of canonical splice junctions, LSTM-base performs comparably to SpliceViNCI for both the dataset. In the prediction of non-canonical splice junctions from the Type-1 dataset, SpliceViNCI shows a minimum improvement of 2% over LSTM-base and a maximum improvement of 28% over SpliceVec-MLP. In the case of the Type-2 dataset, SpliceViNCI shows a maximum and minimum improvement of 2% and 15% over LSTM-base and SpliceRover, respectively. Furthermore, all the models obtain better performance in the identification of non-canonical splice junctions from the Type-2 dataset compared to the Type-1 dataset. This suggests that the negative data in the Type-2 dataset enables the models to learn better representations of the splicing features, as explained in Section 5.2.

### 5.4. Donor and acceptor splicing signals identify the splice junctions cooperatively

We consider donor-acceptor junction pairs as the input sequences instead of only the donor or acceptor junctions. This results in the performance improvement of the predictive model, as shown in Table 4 for the Type-2 dataset. We see that all the models except DeepSplice perform better on the prediction of donor junctions compared to acceptor junctions. However, the performance improves significantly when donor-acceptor junction pairs are considered as input suggesting the cooperative mechanism of donor and acceptor splicing signals in splice junction recognition. This is also inferred in various studies which state that the donor and acceptor junctions are not recognized through individual splicing signals but through junction pairs across exons or introns.[52, 15, 53]

**Table 4:**
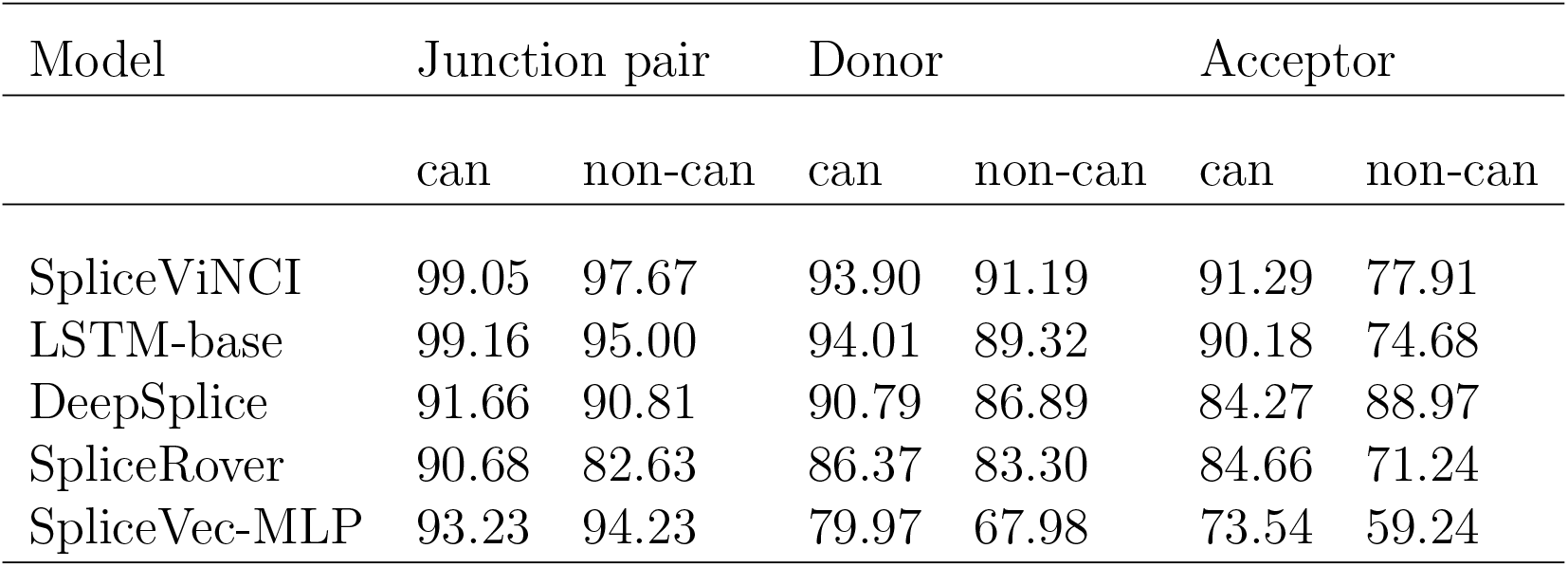
Performance of various state-of-the-art models on Type-2 dataset considering donor, acceptor and donor-acceptor junction pair as input. F1-score (in percentage) is computed as the performance metric.

### 5.5. Identification of novel non-canonical splice junctions by SpliceViNCI

We intend to assess the performance of SpliceViNCI on the identification of novel non-canonical splice junctions, as described in Section 4.1. With this objective, we identify two sets of dimer pairs: seen and unseen. The set of *seen* dimer pairs comprise those positive non-canonical splice junction pairs that are present in both training and test data. Whereas, the set of *unseen* dimer pairs comprise those positive non-canonical splice junctions that are present in test data but not in training data.

Figure 4(a) shows the total number of dimer pairs present in both *seen* and *unseen* categories. The figure also shows the number of dimer pairs correctly identified by SpliceViNCI as splice junctions from *seen* and *unseen* sets in case of both Type-1 and Type-2 dataset. Since the positive data is same in both Type-1 and Type-2 dataset, hence the number of dimer pairs belonging to *seen* and *unseen* sets are same for both the dataset. We observe that SpliceViNCI performs better in the identification of *seen* data in the case of the Type-2 dataset compared to the Type-1 dataset. It is also noteworthy that in case of *unseen* data, SpliceViNCI does not identify any dimer pair from Type-1 dataset, whereas it identifies all except one dimer pairs in case of Type-2 dataset.

**Figure 4:**
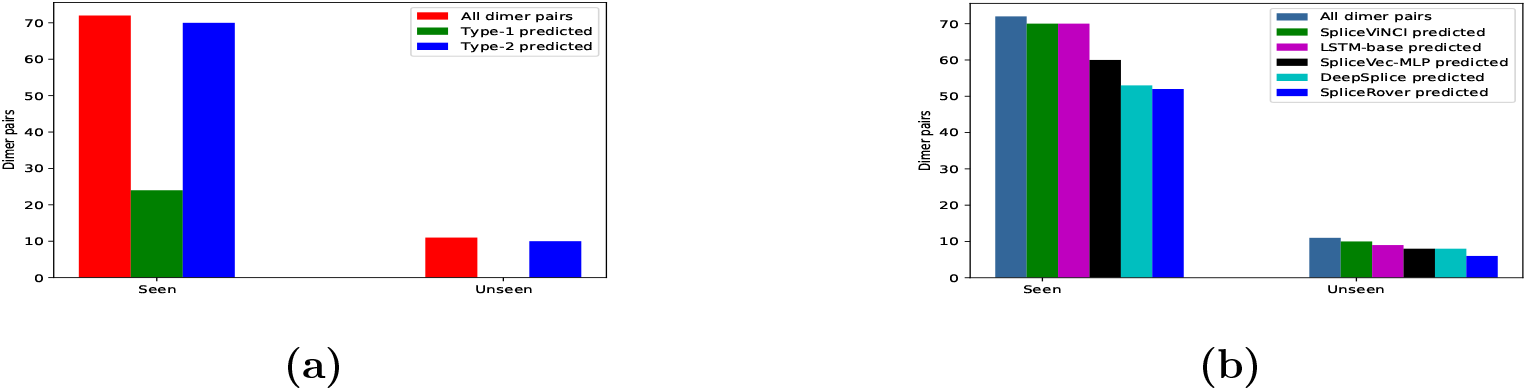
Performance of SpliceViNCI on *seen* and *unseen* data. (a) The number of *seen* and *unseen* dimer pairs identified by SpliceViNCI from Type-1 and Type-2 dataset. (b) The number of *seen* and *unseen* dimer pairs identified by SpliceViNCI and SpliceRover from Type-2 dataset.

We were also curious to observe the performance of all the baselines in the identification of *unseen* data. Figure 4(b) shows the number of dimer pairs identified by SpliceViNCI and all the baselines from both *seen* and *unseen* sets in case of Type-2 dataset. We consider the Type-2 dataset for the analysis because the Type-2 dataset can capture more relevant non-canonical features, as shown in Figure 3 and explained in Section 5.2. It is also fair to compare all the baselines with the Type-2 dataset because we see in Section 5.3 that all the baselines obtain a better performance in the case of the Type-2 dataset. We observe that SpliceViNCI outperforms all the baselines in the identification of both seen and unseen dimer pairs.

### 5.6. Visualization of splicing features captured by SpliceViNCI

The presence of splicing features in the vicinity of splice junctions facilitate the identification of splice junctions by the prediction model. Interpretation of the features captured by the prediction model is necessary to justify the superior performance of the model. The splicing features analyzed using the visualization techniques are summarized in the subsequent subsections. The visualizations are carried out on the non-canonical sequences from the Type-2 dataset. Since our visualizations are specifically for the non-canonical splice junctions, we have considered the Type-2 dataset, which captures more relevant features for the non-canonical splice junctions.

### 5.6.1. Significance of sequence positions

We plot the average *deviation values* obtained for donor and acceptor junction pairs from integrated gradients (Figures 5(a) and 5(c), respectively) and occlusion-1 (Figures 5(b) and 5(d), respectively). A flanking upstream and downstream region of 120 nt is also considered. We observe that both the visualization techniques identify the acceptor and donor splice junctions as the most significant. The importance decreases as the distance of the nucleotide position increases from the splice junction. It is also noteworthy that the importance of the intronic region is higher than the exonic region at both donor and acceptor junctions.

**Figure 5:**
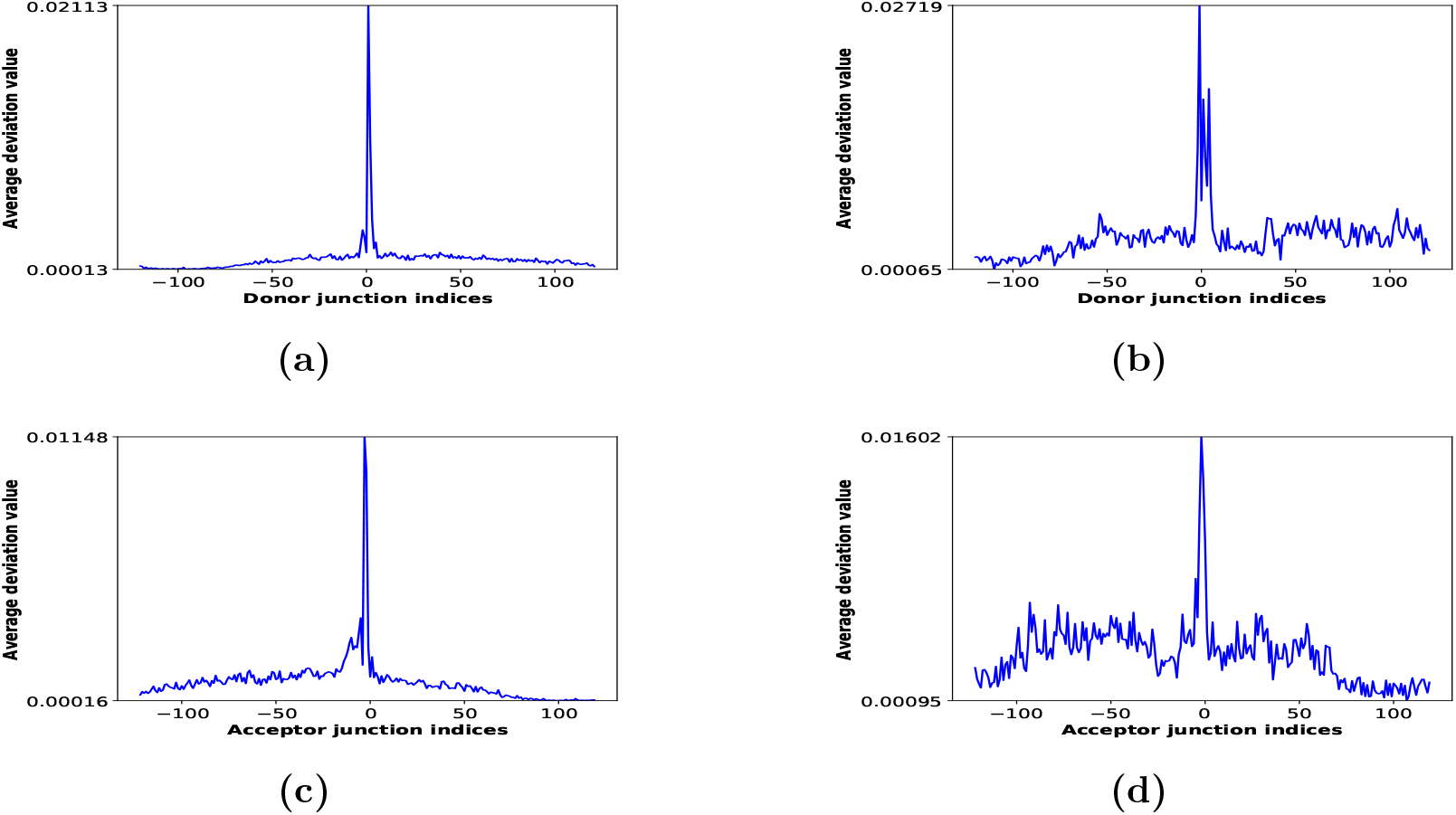
The importance of sequence position. The average deviation value per position is shown for non-canonical donor junctions by (a) integrated gradients (b) occlusion-1 and non-canonical acceptor junctions by (c) integrated gradients (d) occlusion-1.

Additionally, integrated gradients show higher deviation values just upstream of the acceptor junction in the region 0 nt to -15 nt compared to the downstream region, which is due to the presence of weakly conserved polypyrimidine tract (PY-tract) in non-canonical splice junctions.[20] On the other hand, occlusion-1 shows higher importance for sequence positions deeper into the intronic region at the acceptor junction than integrated gradients. This suggests the presence of splicing features upstream of the PY tract.

The higher importance upstream of the PY-tract is captured by occlusion-1 but not integrated gradients. This can be explained with a study by Ancona et al. [40], which states that occlusion is capable of capturing individual features in isolation, whereas integrated gradients perform better when multiple features are considered together. Dutta et al. [19] reported a similar observation where isolated nucleotides were captured as splicing features by occlusion, and consecutive nucleotides were captured as features by integrated gradients. If we consider each sequence position as a feature, then higher *deviation value* assigned by occlusion-1 upstream of the PY-tract suggests the presence of dispersed nucleotides in this region that possibly act as splicing features. This is also stated in a study by Murray et al. [20].

#### 5.6.2. Optimal feature length per position

As the significance of sequence positions observed from Section 5.6.1 suggests the presence of dispersed and isolated splicing features along the intronic region at acceptor junction, we intend to assess the optimal feature length at each sequence position. To this end, we plot the relative frequency of each window length across all sequences when it produced the highest deviation value at a particular sequence position. We observe that at both donor (Figure 6(a)) and acceptor (Figure 6(b)) splice junctions, the plot consistently displays highest frequency for the window length of 1 nt along the entire sequence. This again validates that the features governing non-canonical splicing are mostly dispersed along the sequence and are not placed at consecutive nucleotide positions.

**Figure 6:**
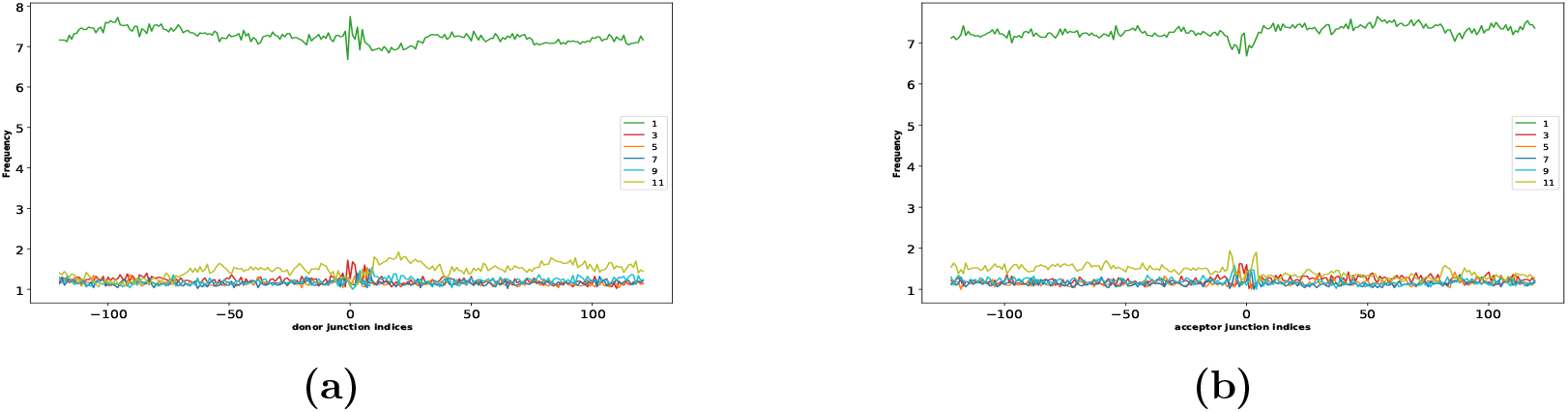
The optimal motif length per position. The frequency of different window lengths, varying from 1 to 11, is shown for occlusion of non-canonical (a) donor and (b) acceptor splice junctions.

#### 5.6.3. Importance of each nucleotide per position

To access the importance of each nucleotide at each sequence position, we plot the average deviation value per position per nucleotide. Integrated gradients show higher deviation value for *C* and *T* along the entire intronic region at both donor (Figure 7(a)) and acceptor junctions (Figure 7(c)). The deviation values for all the nucleotides diminish along the exonic region.

**Figure 7:**
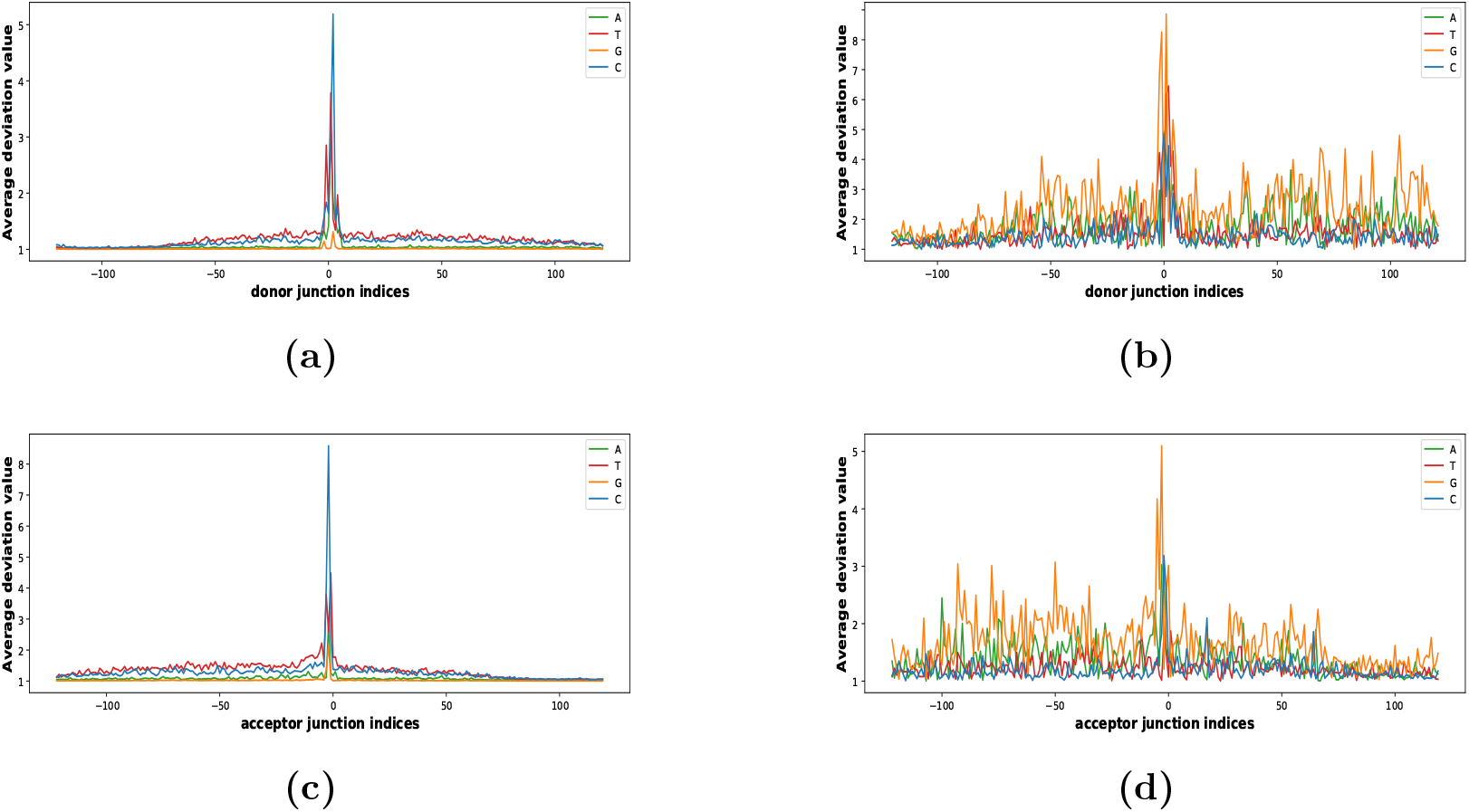
The average deviation value per position per nucleotide. The average deviation value per position per nucleotide is shown for non-canonical donor junctions by (a) integrated gradients (b) occlusion-1 and non-canonical acceptor junctions by (c) integrated gradients (d) occlusion-1.

Occlusion extracts *G* as the most important nucleotide along the intronic region for both donor and acceptor junctions. The deviation value for *G* is particularly high in the region from -30 nt to -100 nt upstream of the acceptor junction, as shown in Figure 7(d). This suggests the presence of a G-rich splicing feature in this region.

The above two observations corroborate the study by Murray et al. [20], which suggests the presence of G-rich motifs upstream of PY-tract in the case of non-canonical introns having weak PY-tract. The integrated gradient captures the weak PY-tract, whereas occlusion captures the G-rich motifs. However, the higher deviation values for *C* and *T* at the donor junction does not relate to any relevant knowledge from the literature.

#### 5.6.4. Most important motifs in a specific region

Murray et al. explored the region -30 nt to -80 nt upstream of the acceptor junction to identify the enriched motifs that regulate splicing in the case of non-canonical splicing. They characterized the relative enrichment of all 4-7 nucleotide k-mers in the specified region and obtained several G-rich motifs.[20] A similar region is also highlighted in Figure 7(d) and described in Section 5.6.3.

We conducted a similar analysis by computing the relative frequency of the most important 4-7 nucleotide k-mers in the region -30 nt to -80 nt relative to the acceptor junction. The most important k-mer in a particular position is the k-mer, which obtains the highest average deviation value, at that position, across all sequences. The motifs obtained for the four different lengths are shown in Figure 8. We observe that the specific region is rich in [AG] nucleotides suggesting the importance of purines in the regulation of non-canonical splicing.

**Figure 8:**
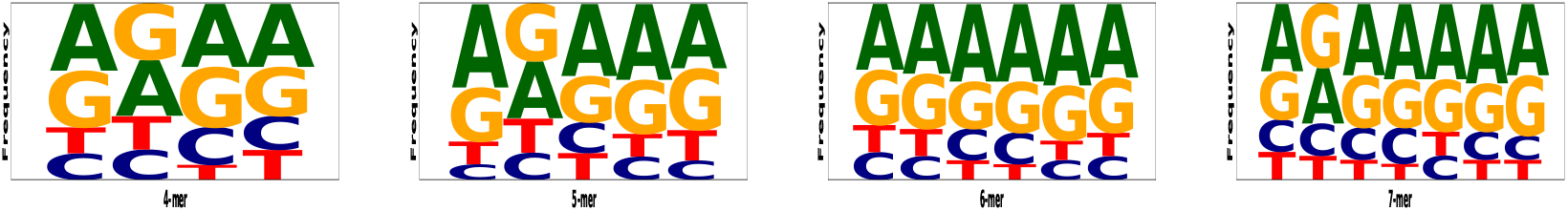
The frequency of various k-mers, given by occlusion, in the region -30 nt to -80 nt upstream of the non-canonical acceptor junction.

To sum up, we can conclude that both the visualization techniques play a vital role in the extraction of non-canonical splicing features. Since the non-canonical splicing features comprise both single and contiguous nucleotides, we can apply both the visualization techniques for the comprehensive under-standing of non-canonical splicing.

## 6. Conclusion

We propose a BLSTM based prediction model named SpliceViNCI that achieves state-of-the-art performance in the identification of canonical and non-canonical splice junctions. SpliceViNCI outperforms state-of-the-art models in the identification of both annotated and novel splice junctions. Our study finds that a flanking region of 120 nt produces optimal performance in the prediction of non-canonical splice junctions.

We employ two effective visualization techniques for the extraction of relevant splicing features learned by the model. The visualization techniques are redesigned to be capable of comprehending genome sequences as input. We employ both back-propagation based and perturbation based visualization techniques to leverage the benefit of both the techniques. The features obtained are validated with the existing knowledge from the literature.

## Acknowledgments

K.K. Singh acknowledges the grant from SERB (CRG/2019/001352). We acknowledge the Department of Biotechnology, Govt. of India for the financial support for the project BT/COE/34/SP28408/2018.

## References

[1] N. Shomron, C. Levy, MicroRNA-biogenesis and pre-mRNA splicing crosstalk, BioMed Research International 2009 (2009).

[2] E. T. Wang, R. Sandberg, S. Luo, I. Khrebtukova, L. Zhang, C. Mayr, S. F. Kingsmore, G. P. Schroth, C. B. Burge, Alternative isoform regulation in human tissue transcriptomes, Nature 456 (2008) 470.

[3] R. Soemedi, K. J. Cygan, C. L. Rhine, J. Wang, C. Bulacan, J. Yang, P. Bayrak-Toydemir, J. McDonald, W. G. Fairbrother, Pathogenic variants that alter protein code often disrupt splicing, Nature genetics 49 (2017) 848.

[4] M. A. Havens, D. M. Duelli, M. L. Hastings, Targeting RNA splicing for disease therapy, Wiley Interdisciplinary Reviews: RNA 4 (2013) 247–266.

[5] M. Aebi, H. Hornig, R. Padgett, J. Reiser, C. Weissmann, Sequence requirements for splicing of higher eukaryotic nuclear pre-mRNA, Cell 47 (1986) 555–565.

[6] M. Kellis, B. Wold, M. P. Snyder, B. E. Bernstein, A. Kundaje, G. K. Marinov, L. D. Ward, E. Birney, G. E. Crawford, J. Dekker, et al., Defining functional dna elements in the human genome, Proceedings of the National Academy of Sciences 111 (2014) 6131–6138.

[7] C. R. Sibley, L. Blazquez, J. Ule, Lessons from non-canonical splicing, Nature Reviews Genetics 17 (2016) 407.

[8] U. Koller, V. Wally, J. W. Bauer, E. M. Murauer, Considerations for a successful rna trans-splicing repair of genetic disorders, Molecular Therapy-Nucleic Acids 3 (2014).

[9] H. Chao, S. G. Mansfield, R. C. Bartel, S. Hiriyanna, L. G. Mitchell, M. A. Garcia-Blanco, C. E. Walsh, Phenotype correction of hemophilia a mice by spliceosome-mediated rna trans-splicing, Nature medicine 9 (2003) 1015–1019.

[10] Y. Li, C.-y. Chen, A. M. Kaye, W. W. Wasserman, The identification of cis-regulatory elements: A review from a machine learning perspective, Biosystems 138 (2015) 6–17.

[11] S. Min, B. Lee, S. Yoon, Deep learning in bioinformatics, Briefings in bioinformatics 18 (2017) 851–869.

[12] Y. Zhang, X. Liu, J. MacLeod, J. Liu, Discerning novel splice junctions derived from RNA-seq alignment: a deep learning approach, BMC genomics 19 (2018) 971.

[13] J. Zuallaert, F. Godin, M. Kim, A. Soete, Y. Saeys, W. De Neve, SpliceRover: interpretable convolutional neural networks for improved splice site prediction, Bioinformatics 34 (2018) 4180–4188.

[14] B. Lee, T. Lee, B. Na, S. Yoon, DNA-level splice junction prediction using deep recurrent neural networks, arXiv preprint arXiv:1512.05135 (2015)

[15] K. Jaganathan, S. K. Panagiotopoulou, J. F. McRae, S. F. Darbandi, D. Knowles, Y. I. Li, J. A. Kosmicki, J. Arbelaez, W. Cui, G. B. Schwartz, et al., Predicting splicing from primary sequence with deep learning, Cell 176 (2019) 535–548.

[16] T. Lee, S. Yoon, Boosted categorical restricted boltzmann machine for computational prediction of splice junctions, in: International Conference on Machine Learning, 2015, pp. 2483–2492.

[17] Y. Zhang, X. Liu, J. N. MacLeod, J. Liu, DeepSplice: Deep classification of novel splice junctions revealed by RNA-seq, in: 2016 IEEE International Conference on Bioinformatics and Biomedicine (BIBM), IEEE, 2016, pp. 330–333.

[18] A. Dutta, T. Dubey, K. K. Singh, A. Anand, SpliceVec: distributed feature representations for splice junction prediction, Computational biology and chemistry 74 (2018) 434–441.

[19] A. Dutta, A. Dalmia, R. Athul, K. K. Singh, A. Anand, Using the chou’s 5-steps rule to predict splice junctions with interpretable bidirectional long short-term memory networks, Computers in Biology and Medicine (2019).

[20] J. I. Murray, R. B. Voelker, K. L. Henscheid, M. B. Warf, J. A. Berglund, Identification of motifs that function in the splicing of non-canonical introns, Genome biology 9 (2008) R97.

[21] C. Trapnell, L. Pachter, S. L. Salzberg, TopHat: discovering splice junctions with RNA-seq, Bioinformatics 25 (2009) 1105–1111.

[22] K. F. Au, H. Jiang, L. Lin, Y. Xing, W. H. Wong, Detection of splice junctions from paired-end rna-seq data by SpliceMap, Nucleic acids research 38 (2010) 4570–4578.

[23] M. Pertea, X. Lin, S. L. Salzberg, GeneSplicer: a new computational method for splice site prediction, Nucleic acids research 29 (2001) 1185–1190.

[24] S. Degroeve, Y. Saeys, B. De Baets, P. Rouzé, Y. Van de Peer, SpliceMachine: predicting splice sites from high-dimensional local context representations, Bioinformatics 21 (2004) 1332–1338.

[25] J. Huang, T. Li, K. Chen, J. Wu, An approach of encoding for prediction of splice sites using SVM, Biochimie 88 (2006) 923–929.

[26] A. K. Baten, B. C. Chang, S. K. Halgamuge, J. Li, Splice site identification using probabilistic parameters and SVM classification, BMC bioinformatics 7 (2006) S15.

[27] S. Sonnenburg, G. Schweikert, P. Philips, J. Behr, G. Rätsch, Accurate splice site prediction using support vector machines, BMC bioinformatics 8 (2007) S7.

[28] S. Degroeve, B. De Baets, Y. Van de Peer, P. Rouzé, Feature subset selection for splice site prediction, Bioinformatics 18 (2002) S75–S83.

[29] Y. Saeys, S. Degroeve, Y. Van de Peer, Digging into acceptor splice site prediction: an iterative feature selection approach, in: European Conference on Principles of Data Mining and Knowledge Discovery, Springer, 2004, pp. 386–397.

[30] M. G. Reese, F. H. Eeckman, D. Kulp, D. Haussler, Improved splice site detection in genie, Journal of computational biology 4 (1997) 311–323.

[31] M. Du, N. Liu, X. Hu, Techniques for interpretable machine learning, Communications of the ACM 63 (2019) 68–77.

[32] A. Shrikumar, P. Greenside, A. Kundaje, Learning important features through propagating activation differences, in: Proceedings of the 34th International Conference on Machine Learning-Volume 70, JMLR. org, 2017, pp. 3145–3153.

[33] S. Hochreiter, J. Schmidhuber, Long short-term memory, Neural computation 9 (1997) 1735–1780.

[34] F. A. Gers, J. Schmidhuber, F. Cummins, Learning to forget: Continual prediction with LSTM (1999).

[35] F. A. Gers, N. N. Schraudolph, J. Schmidhuber, Learning precise timing with LSTM recurrent networks, Journal of machine learning research 3 (2002) 115–143.

[36] A. Santoro, R. Faulkner, D. Raposo, J. Rae, M. Chrzanowski, T. Weber, D. Wierstra, O. Vinyals, R. Pascanu, T. Lillicrap, Relational recurrent neural networks, in: Advances in Neural Information Processing Systems, 2018, pp. 7299–7310.

[37] D. P. Kingma, J. Ba, Adam: A method for stochastic optimization, ICLR (2015).

[38] M. Sundararajan, A. Taly, Q. Yan, Axiomatic attribution for deep networks, in: Proceedings of the 34th International Conference on Machine Learning-Volume 70, JMLR. org, 2017, pp. 3319–3328.

[39] A. Binder, G. Montavon, S. Lapuschkin, K.-R. Müller, W. Samek, Layer-wise relevance propagation for neural networks with local renormalization layers, in: International Conference on Artificial Neural Networks, Springer, 2016, pp. 63–71.

[40] M. Ancona, E. Ceolini, C. Oztireli, M. Gross, Towards better understanding of gradient-based attribution methods for deep neural networks, in: 6th International Conference on Learning Representations (ICLR 2018), 2018.

[41] N. Poerner, H. Schütze, B. Roth, Evaluating neural network explanation methods using hybrid documents and morphological prediction, in: 56th Annual Meeting of the Association for Computational Linguistics (ACL), 2018.

[42] S. M. Lundberg, S.-I. Lee, A unified approach to interpreting model predictions, in: Advances in neural information processing systems, 2017, pp. 4765–4774.

[43] M. D. Zeiler, R. Fergus, Visualizing and understanding convolutional networks, in: European conference on computer vision, Springer, 2014, pp. 818–833.

[44] J. Harrow, A. Frankish, J. M. Gonzalez, E. Tapanari, M. Diekhans, F. Kokocinski, B. L. Aken, D. Barrell, A. Zadissa, S. Searle, et al., GENCODE: the reference human genome annotation for the ENCODE project, Genome research 22 (2012) 1760–1774.

[45] A. Piovesan, M. Caracausi, M. Ricci, P. Strippoli, L. Vitale, M. C. Pelleri, Identification of minimal eukaryotic introns through geneBase, a user-friendly tool for parsing the NCBI gene databank, DNA Research 22 (2015) 495–503.

[46] M. O. Noordewier, G. G. Towell, J. W. Shavlik, Training knowledge-based neural networks to recognize genes in DNA sequences, in: Advances in neural information processing systems, 1991, pp. 530–536.

[47] M. Burset, I. Seledtsov, V. Solovyev, Analysis of canonical and non-canonical splice sites in mammalian genomes, Nucleic acids research 28 (2000) 4364–4375.

[48] S. M. Mount, Genomic sequence, splicing, and gene annotation, The American Journal of Human Genetics 67 (2000) 788–792.

[49] A. Bari, M. R. Reaz, B.-S. Jeong, Effective dna encoding for splice site prediction using SVM, MATCH Commun. Math. Comput. Chem 71 (2014) 241–258.

[50] L. v. d. Maaten, G. Hinton, Visualizing data using t-SNE, Journal of machine learning research 9 (2008) 2579–2605.

[51] H. Bretschneider, S. Gandhi, A. G. Deshwar, K. Zuberi, B. J. Frey, Cossmo: predicting competitive alternative splice site selection using deep learning, Bioinformatics 34 (2018) i429–i437.

[52] S. M. Berget, Exon recognition in vertebrate splicing, Journal of biological Chemistry 270 (1995) 2411– 2414.

[53] H. Shenasa, K. J. Hertel, Combinatorial regulation of alternative splicing, Biochimica et Biophysica Acta (BBA)-Gene Regulatory Mechanisms (2019).

